# Integrating brain methylome with GWAS for psychiatric risk gene discovery

**DOI:** 10.1101/440206

**Authors:** Shizhong Han, Ying Lin, Minghui Wang, Fernando S. Goes, Kai Tan, Peter Zandi, Thomas Hyde, Daniel R. Weinberger, James B. Potash, Joel E. Kleinman, Andrew E. Jaffe

## Abstract

DNA methylation (DNAm) is heritable and plays a role in brain development and function through transcriptional regulation. Aberrant DNAm in human brain has been linked to psychiatric disorders, potentially as mediators of common genetic risk variants. In this study, we hypothesize that common risk variants for psychiatric disorders may act through affecting DNAm level in human brain. We first aimed to investigate the heritability pattern of DNAm levels in the human prefrontal cortex. Secondly, through imputation-driven methylome-wide association study (MWAS), we aimed to identify CpG sites whose methylation levels are genetically associated and that show methylation-trait associations in the prefrontal cortex of patients with schizophrenia or bipolar disorder. Our heritability analysis showed that, of ~370,000 CpG sites measured with the Illumina HumanMethylation450 microarray, 17% were heritable (p < 0.05), with a mean heritability of 0.22. Heritable CpG sites were enriched in intergenic regions, CpG shore, and regulatory regions in prefrontal cortex. Our MWAS approach identified known and potentially novel risk genes harboring CpG sites of methylation-trait associations for schizophrenia or bipolar disorder, which were not detectable using three alternative strategies (blood-based methylome reference, transcriptome-wide association study, and two gene-based association tests). Gene set enrichment analysis for genes with methylation-trait association evidence revealed pathways clearly related to neuronal functions, but also highlighted additional biological mechanisms that may underlie psychiatric disorders, such as microRNA-related regulation. In conclusion, our results showed the power of integrating brain methylation data with GWAS for psychiatric risk gene discovery, with potential applications in brain-related disorders or traits.

## Introduction

DNA methylation (DNAm) is heritable and plays a critical role in brain development and function through transcriptional regulation (1, 2). Family and twin studies have investigated the heritability of DNAm for sites across the genome in easily accessible tissues (3–7), but, to our knowledge, there have been limited studies of methylation heritability in brain tissue. Indeed, only one study estimated the heritability of DNAm levels of individual CpG dinucleotides attributable to local single nucleotide polymorphisms (SNPs) in postmortem brain from unrelated individuals, but the estimation was limited to ~21,000 CpG sites primarily within promoters (8). The heritability pattern is not known for DNAm sites within gene body or intergenic regions, which represent a large portion of DNAm variation and are potentially important in epigenetic regulation of gene expression (9–12).

Aberrant DNAm has been linked to psychiatric disorders in candidate gene analysis and epigenome-wide association studies (EWAS) of schizophrenia (SCZ) (13, 14), bipolar disorder (BD) (15), and major depressive disorder (16, 17). However, the majority of these studies have been of easily accessible peripheral tissue (blood), which may not correlate with methylation signals in the brain (18, 19). Additionally, due to the challenges of accessing brain tissues, studies that examined brain samples were often limited by small sample size. Finally, it is hard to establish a causal role between DNAm change and disease as it is unclear whether aberrant DNAm causes disease or vice versa.

Genome-wide association studies (GWAS) indicate that most common genetic risk variants underlying complex diseases are noncoding and enriched in regulatory genomic regions in cells or tissues related to disease (20), suggesting that risk variants may act through regulation of cell/tissue-specific molecular mechanisms, such as methylation or gene expression. For example, SCZ risk variants from GWAS were enriched at enhancers active in brain, but not in tissues unlikely to be relevant to SCZ (21). Furthermore, statistical genetic approaches like linkage disequilibrium (LD) score regression have shown enrichment of trait heritability in regions with functional annotations that are trait-related and cell-type-specific, including enrichment of heritability for SCZ and BP in functional regions specific to the central nervous system (22) and even specific subclasses of neurons (23).

In line with a heritable component for DNAm, studies of methylation quantitaive trait loci (meQTL) have identified specific SNPs affecting methylation levels of CpG sites in human cell lines (24, 25), peripheral tissues (26, 27), and the human brain (13, 28, 29). SNPs associated with meQTL largely act in *cis* and tend more often to be located at distant regulatory regions than at promoters (30, 31). Jaffe et al have shown that a large fraction of GWAS risk loci for SCZ contains an meQTL detected in adult frontal cortex (13). Significant enrichment of SCZ risk variants has also been observed among meQTLs calculated in prenatal brain tissue (29). It is therefore likely that DNAm may mediate the effect of genotype on disease risk for a proportion of risk variants.

In the current study, we first aimed to investigate the extent to which DNAm levels are determined by cis-genetic variations in human prefrontal cortex for ~370,000 CpG sites measured with the Illumina HumanMethylation450 microarray. Secondly, through imputation of methylation-trait association statistics from GWAS summary statistics, we aimed to identify CpG sites whose methylation is genetically associated, and that show methylation-trait associations in the prefrontal cortex for two major psychiatric disorders ‒ SCZ and BD.

## Methods

### Datasets

DNAm and GWAS SNP genotyping data were available from postmortem dorsolateral prefrontal cortex brain tissue samples from 526 individuals. Methylation was measured using the Illumina HumanMethylation450 (450k) microarray, which measures CpG methylation across >485,000 probes. The current study used DNAm data from 238 unrelated subjects of European ancestry (100 SCZ patients and 138 controls), defined by principal component analysis (PCA) of GWAS data collected from the same samples. DNAm data were processed and normalized using the Minfi Bioconductor package in R (32). GWAS data were imputed into 1000 Genomes Phase 3 variants using SHAPEIT2 (33) and IMPUTE2 (34). After filtering out SNPs with MAF < 5%, HWE p-value < 0.05, or missing rate > 10%, there were 4,402,285 SNPs on autosomes retained for further analyses. Information on tissue processing, experimental and bioinformatics procedures related to the methylation data, and genotype data processing was described in prior reports (1, 13).

We downloaded GWAS summary statistics from the Psychiatric Genomic Consortium (PGC) website (https://www.med.unc.edu/pgc/) for SCZ and BD. We then applied an imputation-driven methylome-wide association study (MWAS) to the PGC2 GWAS meta-analysis summary statistics. We retained autosomal SNPs with MAF > 0.05 and imputed quality score > 0.8. Coordinates of SNPs were aligned to hg19 for all datasets.

### SNP-heritability of DNAm

We estimated SNP-heritability of DNAm for CpG sites using GCTA software (35), which calculates the genetic relationship matrix (GRM) between subjects and uses the restricted maximum likelihood to estimate variance components in a mixed model framework. All pairs of 238 subjects with both GWAS and methylation data had a genetic relatedness 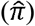 less than 0.025 and were included for heritability estimation. GRM was calculated using SNPs within 1Mb of each CpG site. A number of covariates were included in the mixed model, including age, sex, SCZ diagnosis, the top five principal components (PCs) from the GWAS data to account for ancestry variability, and the top ten PCs from the methylation data (beta values) to control for cellular heterogeneity and latent batch effects. To avoid technical noise arising from SNPs within a probe region, we limited heritability estimation to 370,538 CpG sites whose probes did not overlap with any SNPs based on annotations in the *minfi* Bioconductor package in R (32).

### CpG annotations

To investigate the distribution of heritable CpG sites (p < 0.05) across genomic features, we annotated CpG sites within three different contexts: 1) gene-based annotations from ANNOVAR (36), including exonic, splicing, ncRNA, 5’UTR, 3’UTR, intron, upstream (1 kb upstream of transcription start site), downstream (1 kb upstream of transcription end site), and intergenic; 2) annotations by distance to CpG island, including island, shelf (~4 kb from islands), shore (~2 kb from islands), and sea (others) using annotation information from the Illumina Infinium HM450 manifest file; 3) functional annotations of 15-core chromatin states, derived using hidden Markov models, in the dorsolateral prefrontal cortex (E073) sample from the Epigenome Roadmap project (37). We used simulation-based approach to estimate the enrichment statistics of heritable CpG sites in each annotation category. Specifically, we first generated a background distribution from random CpG sets, while matching for CpG density found in heritable CpG sites. The enrichment fold was estimated by the ratio of the observed number of heritable CpGs overlapping with a specific annotation category to the average number of that from random CpG sets. The p-value for enrichment (depletion) was then the proportion of random CpG sets that fall in the same or a greater (smaller) number of annotations in each annotation category, as compared to the number found within heritable CpG set.

### Estimation of the *cis*-genetic component of methylation

For each heritable CpG site (p < 0.05), we first corrected its methylation level (beta value) by regressing out the same set of covariates as included in the heritability analysis. We then built models to predict the residual methylation levels using SNPs within 1Mb of each CpG. We investigated three modeling schemes: top SNP, polygenic score, and elastic net. In the top SNP model, the SNP was simply chosen as the one with the strongest association with methylation. In the polygenic score model, we included all SNPs that were associated with methylation level at a significance level of p < 0.05, and the weights for the polygenic score were the effect sizes (beta coefficients from regression) estimated from the training sample. In the elastic net model, SNPs were selected using a penalized regression procedure that combines LASSO and ridge regression. The parameter α was tuned from 0 to 1 with a step increase by 0.1. The final α value was determined as the one with the smallest prediction error from tenfold cross validation. To determine the optimal modeling scheme, we compared the fivefold cross-validated prediction *R*^2^ values (the square of the correlation between predicted and observed methylation) for the three modelling schemes.

### Imputation-driven methylome-wide association study (MWAS)

We performed MWAS to identify CpGs sites with methylation-trait association evidence by imputing GWAS summary statistics into methylation-trait association statistics for two psychiatric disorders SCZ and BD. To impute as many CpG sites as possible, we reintroduced CpG sites whose probes overlapped SNPs. Imputation was done for 49,442 CpG sites that were heritable (p < 0.05) and had a cross-validated prediction *R*^2^ > 0.01 by elastic net. Following the same theoretical framework as a transcriptome-wide association study (TWAS) (38), MWAS uses methylation data to compute weights for SNP association statistics from GWAS. In brief, based on our existing brain samples with both methylome and GWAS data, we used elastic net to estimate the genetic effects on DNAm level for SNPs within 1 Mb of each CpG site. The methylation-trait association statistic was then calculated as a weighted linear combination of SNP-trait association statistics from GWAS, with weights being equal to estimated effect sizes on DNAm level from elastic net. Formally, let Z be a vector of the Z-statistic for association between the GWAS trait and SNPs within 1Mb of a CpG site. We define W as a vector of weights that are effect sizes of the same set of SNPs on methylation level derived by elastic net. The imputed Z-statistic of methylation-trait association is then *WZ*/(*W* Σ *W*)^1/2^, where Σ is the correlation matrix among SNPs that can be estimated from an ancestry-matched reference sample. Bonferroni correction was used to control for multiple testing of 49,442 CpGs with a significance threshold of p = 1×10^−6^ for each disorder. We made the SNP weights on DNAm for each CpG site publicly available at: https://figshare.com/s/052a0b729c3c7ad7b535.

### TWAS and gene-based association tests

We examined whether significant genes discovered by MWAS could be similarly uncovered by three alternative strategies. The first strategy utilized the MWAS framework, but used SNP weights derived from a blood-based methylome reference from http://mcn.unibas.ch/files/EstiMethDistribution.zip (39). The second strategy was TWAS with the weights derived from gene expression data using dorsolateral prefrontal cortex, downloaded from the TWAS website (http://gusevlab.org/projects/fusion/#reference-functional-data). The third strategy involved two gene-level association tests for common variants, GATES (40) and VEGAS-sum (41). We assigned a SNP to a gene if it was located within the gene, based on NCBI 37.3 gene annotation, or within 20 kb upstream or downstream of the gene, to capture cis regulatory variants. GATES uses an extended Simes procedure to derive a gene-based p-value and is powerful when there is only one or a few causal SNPs. VEGAS-sum computes a gene-level p-value using simulations from a multivariate normal distribution and is powerful when there are multiple independent causal SNPs. Pairwise SNP correlations within a gene were estimated based on the genotype data of unrelated CEU samples from the 1000 Genomes Project. The total number of tests were 86,431 (CpGs), 5,419 (genes), and 24,583 (genes) for MWAS-blood, TWAS, and two gene-based tests, respectively. The Bonferroni–corrected significance threshold for each strategy was set at p = 5.8×10^−7^ (MWAS-blood), p = 9×10^−6^ (TWAS), and p = 2×10^−6^ (GATES and VEGAS-sum).

### Gene set enrichment analysis

To understand the biology of genes with CpG sites of methylation-trait associations, we conducted gene set enrichment analysis using the logistic regression-based method LRpath (42), which relates the odds of gene set membership with the gene-level association signals, while adjusting for potentially confounding factors. Specifically, for each gene set, we defined the dependent variable y as 1 for genes in the set, and 0 for all other genes. The independent variable x is −log (p), where p is the gene-based p value defined by the smallest p-value among all imputed methylation-traits associations for CpG sites within a gene. Logistic regression is used to model the log-odds of a gene belonging to a specific gene set as a function of x. We included the number of imputed CpG sites within a gene as a covariate to control for potential confounding effect due to gene size difference. Gene sets were from two resources: 1) gene sets that are potentially important in psychiatric disorders (72 gene sets) (43), and 2) canonical pathways, Gene Ontology, and microRNA targets from MSigDB (v5.2) (5,986 gene sets) (44). To facilitate interpretation of the results, we included gene sets that overlapped at least 20, but not more than 2,000 genes with our tested genes. False discovery rate (FDR) q-values were calculated to account for multiple testing using the R statistical package “qvalue” (www.r-project.org).

We used hierarchical clustering to group significant gene sets into clusters based on similarity of their gene profiles (45). We first defined a gene overlapping matrix by counting the number of overlapping genes for each pair of gene sets. The Pearson correlation coefficient *R* was then calculated for each pair of gene sets based on their overlap profiles. The distance matrix for hierarchical clustering was then 1 − *R*. Hierarchical clustering was performed using the “ward” method implemented in R function “hclust”. The dendrogram and heatmap were plotted using the R function “heatmap.2”. Clusters of related gene sets were manually determined to represent biologically relevant groups.

## Results

### SNP-heritability of DNAm

We estimated the SNP-heritability of DNAm levels for each CpG site using SNPs within 1 Mb. Figure 1a shows the distribution of heritability estimates for all, heritable (p ≤ 0.05), and non-heritable (p > 0.05) CpG sites separately. The average heritability was 0.050 across all CpG sites, heritable and nonheritable, ranging from 0 to 0.99. There were 62,387 (17%) CpG sites reaching a nominal level of significance for the heritability estimate (p < 0.05), with an average heritability of 0.22, ranging from 0.022 to 0.99. Heritable CpG sites tended to be more continuous (Figure 1b) and had higher variance on average in methylation levels (beta-value) than non-heritable CpG sites (Wilcoxon rank sum test, p < 2.2× 10^−16^) (Figure 1c). We investigated the distribution of heritable CpG sites across genomic features within different contexts. Heritable CpG sites were enriched in intergenic regions, CpG shore, and regulatory genomic regions (flanking active TSS and enhancers) in prefrontal cortex, but were depleted in 5’UTR, CpG islands, active TSS and strong transcription states in prefrontal cortex (Figure 1d).

**Figure 1.**
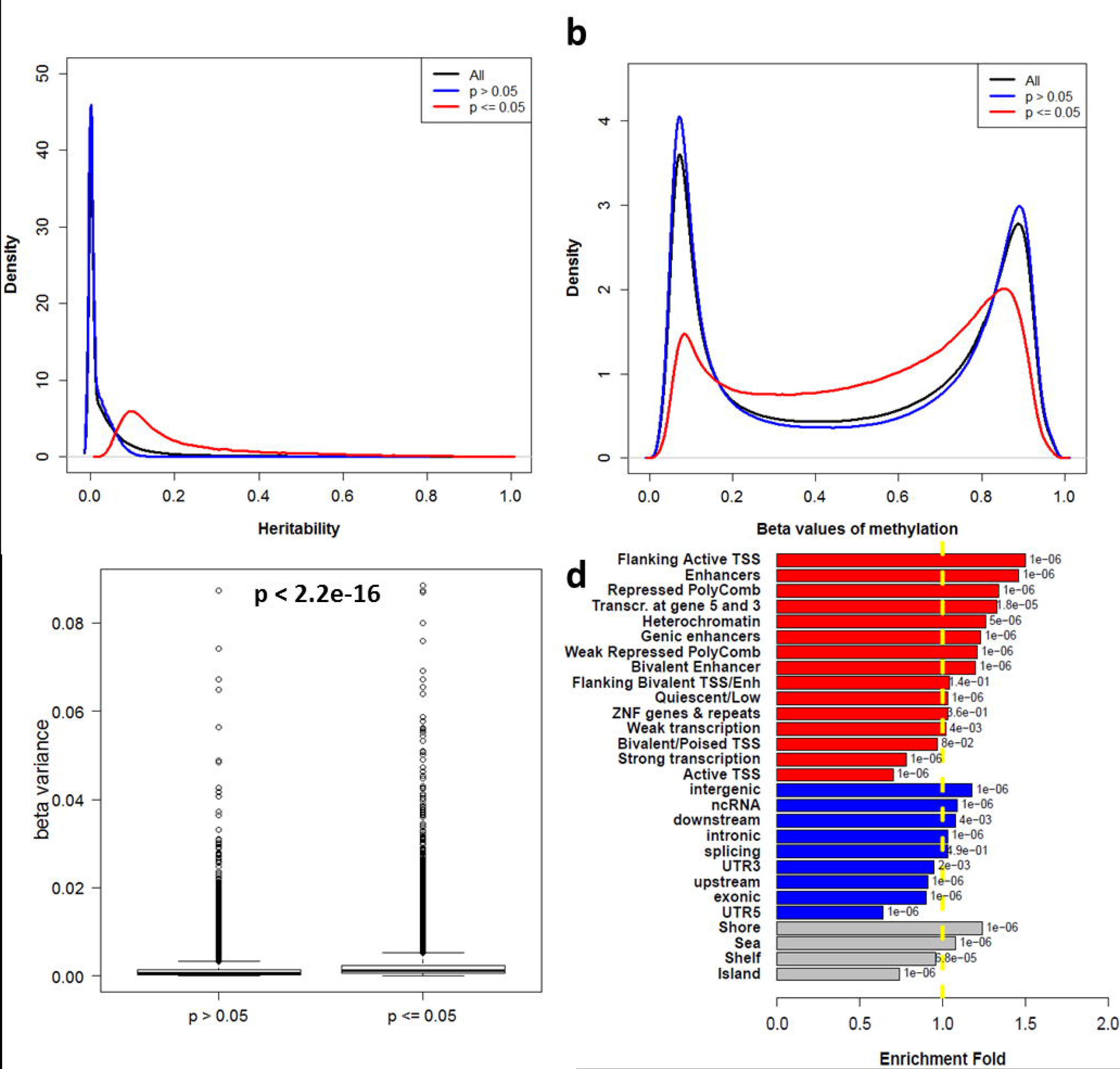
SNP-based heritability analysis of DNAm levels: a) Distribution of heritability estimates for all, heritable (p ≤ 0.05), and non-heritable (p > 0.05) CpG sites; b) Distribution of methylation levels (beta-value) for all, heritable (p ≤ 0.05), and non-heritable (p > 0.05) CpG sites; c) Boxplot for variance of methylation levels (beta-value) for heritable (p ≤ 0.05) and non-heritable (p > 0.05) CpG sites; d) Enrichment of heritable CpG sites across genomic features within three different contexts: genomic locations (blue), distance to CpG islands (grey), and functional states in the dorsolateral prefrontal cortex (red). The x-axis represents enrichment fold. The y-axis labels the different features examined. The dotted yellow line indicates no enrichment (fold = 1). The numbers next to each bar are enrichment p-values.

We further estimated the SNP-heritability of DNAm levels using 100 neurotypical control samples. The heritability estimates were highly correlated between all samples and the neurotypical only samples (*R*^2^ = 85, Supplementary Figure 1). The average heritability in the neurotypical only sample was 0.062 across all CpG sites, which was slightly higher than the average estimates from the overall samples (0.05). Compared to the overall samples, there were fewer CpGs (48,359, 13%) reaching nominal significance for the heritability estimates (p < 0.05), possibly due to the smaller sample size. In accordance with the pattern observed in the overall samples, heritable CpG sites tended to be more continuous and variable in methylation levels than non-heritable CpG sites, enriched in intergenic regions, CpG shore, flanking active TSS states in prefrontal cortex; heritable CpG sites were depleted in 5’UTR, CpG islands, and active TSS states in prefrontal cortex (Supplementary Figure 2).

### Predicting the *cis*-genetic component of DNAm

We evaluated whether the DNAm levels of heritable CpGs could be imputed from the genotype data for SNPs within 1 Mb of the CpG. We used fivefold cross-validation to compare predictive performance of three modeling schemes: top SNP, polygenic score and elastic net. Of the 32,949 CpGs for which all three models were successfully built, we found that the elastic net approach achieved the best performance with an average *R*^2^=0.24, followed by the polygenic score (*R*^2^=0.16), and the top SNP methods (*R*^2^=0.095) (Figure 2a). When models were compared on the same CpG sites, 80% of the CpG sites achieved the best prediction by elastic net, whereas 18% and 2% of CpG sites were best predicted by the polygenic score and the top SNP methods, respectively (Figure 2b). Using heritability estimates as a benchmark for prediction *R*^2^, we compared the prediction *R*^2^ with the heritability estimate for each CpG (Figure 2c). We found that 89% of the CpGs achieved an *R*^2^ equal to or larger than the lower bound of heritability estimate by elastic net, whereas 68% (polygenic score) and 39% (top SNP) of CpGs achieved such performance with the other methods. Overall, the elastic net approach showed the best performance for predicting DNAm levels based on different evaluation strategies. The effect sizes derived from elastic net were then used to impute differential methylation statistics from GWAS summary statistics.

**Figure 2.**
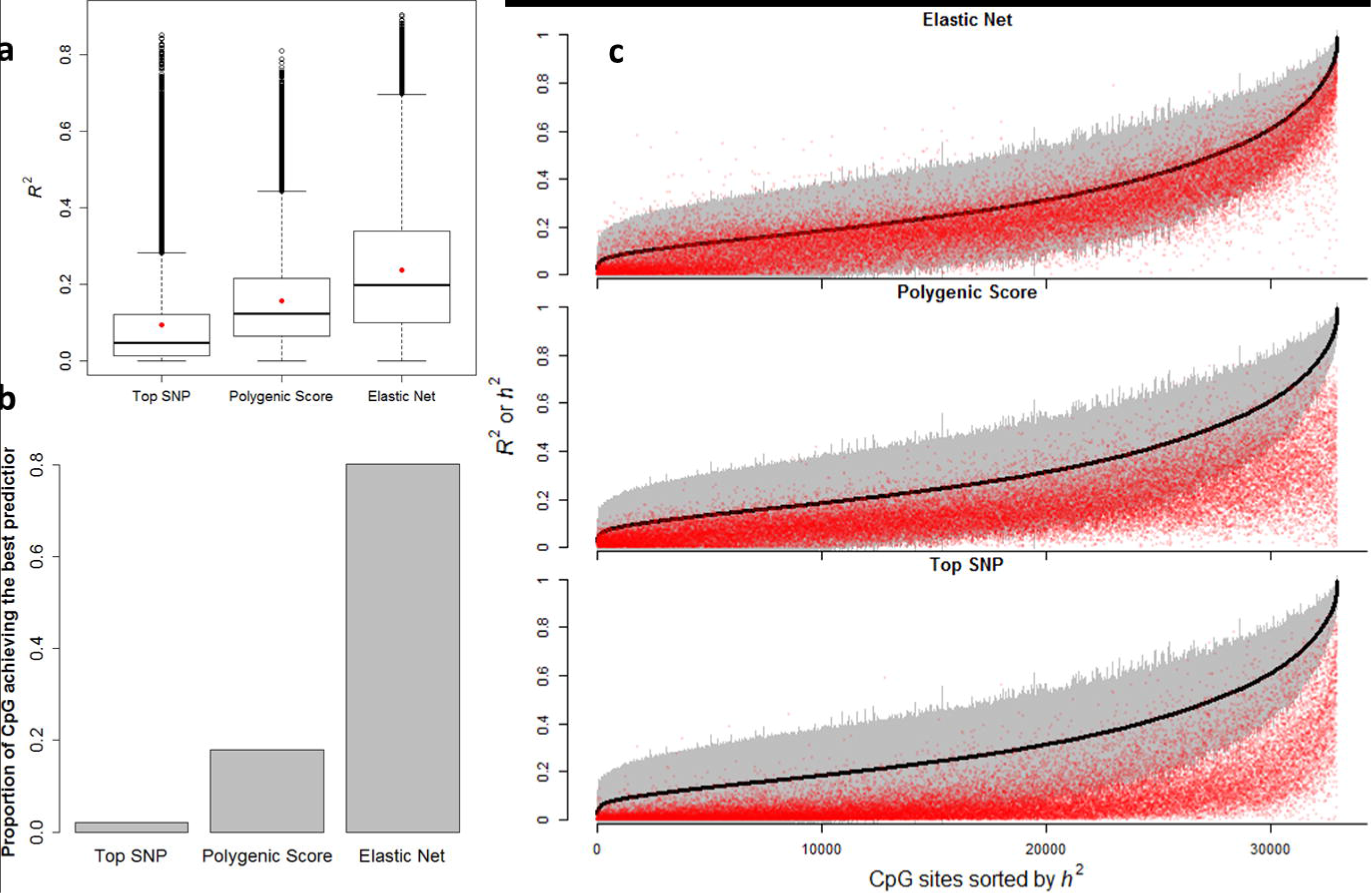
Evaluation of prediction performance (*R*^2^) for three modeling schemes: a) Boxplot of prediction performance (*R*^2^) for three modeling schemes. Red points represent the mean value of *R*^2^ across CpG sites; b) Barplot for the proportion of CpGs achieving the best prediction performance under each modeling scheme; c) Comparison of prediction performance (*R*^2^) with heritability estimates (*h*^2^). The figure shows the cross-validated prediction performance (*R*^2^ in red) in comparison to methylation heritability estimates (black). The gray zone shows the 95% confidence intervals of heritability estimates.

### Imputation-driven methylome-wide association study (MWAS)

We performed imputation-driven MWAS to identify CpGs sites of methylation-trait associations in prefrontal cortex for SCZ and BD. We imputed methylation-trait association statistics from PGC2 GWAS summary statistics for both disorders.

We identified 914 genome-wide significant (GWS) CpGs with differential methylation evidence for SCZ (Figure 3a, Supplementary Table 1). Of these, 883 overlapped original GWAS risk loci (defined by 1 Mb upstream and downstream of any GWS SNPs from PGC2 SCZ GWAS); 677 CpGs were within the extended major histocompatibility complex region (chr6:25652464-33771788); 31 CpG sites were located more than 1 Mb away from any GWS SNPs from PGC2 SCZ GWAS, suggesting potential novel signals not detected by GWAS. Table 1 shows details and methylation-trait association statistics for the 20 CpG sites mapped to 24 genes. Of note, the gene *MORC2-AS1* was far from GWS in PGC2 SCZ GWAS (smallest p = 1.5×10^−5^, 1 Mb window), but contained a CpG site at a gene body with strong evidence for methylation-trait association (cg13896476, p = 3.4×10^−10^). Another CpG site, at promoter of *MORC2-AS1*, also showed significant evidence for methylation-trait association (cg08837037, p = 1.5×10^−8^, Supplementary Table 1). Figure 4 shows the regional association plot of methylation-trait associations around *MORC2-AS1*, along with SNP association signals from GWAS and how their weights contribute to the top significant CpG.

**Figure 3.**
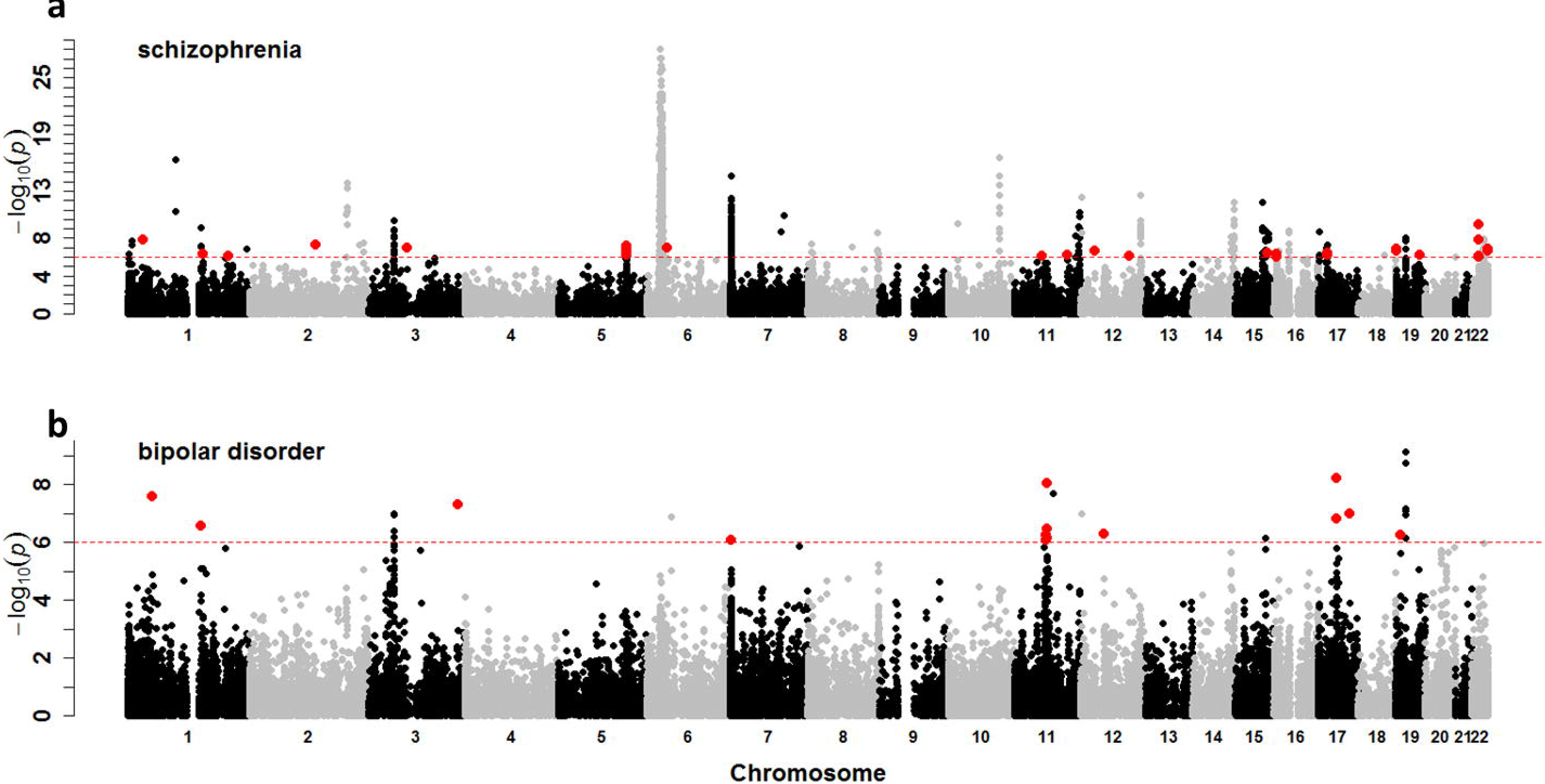
Genome-wide association plot for methylation-trait associations for schizophrenia and bipolar disorder (PGC2). The dotted red line indicates significance thresholds. Red points represent significant CpG sites that are at least 1Mb away from genomewide significant SNPs.

**Table 1.** Information and methylation-trait association statistics for CpGs associated with schizophrenia.

| **CpGs** | **Chr** | **Position** | **Gene** | **Function** | **z** | **p** | **P<sub>smallest</sub><sup>\*</sup>** |
| --- | --- | --- | --- | --- | --- | --- | --- |
| ch.1.1029351R | 1 | 31734918 | *SNRNP40* | Body | -5.66 | 1.50E-08 | 3.51E-06 |
| cg06221963 | 1 | 154839813 | *KCNN3* | Body | -5.01 | 5.31E-07 | 3.18E-06 |
| cg24077277 | 1 | 205201540 | *TMCC2* | Body | -4.95 | 7.28E-07 | 8.69E-07 |
| cg19875535 | 5 | 140030758 | *IK* | Body | -5.40 | 6.51E-08 | 4.28E-07 |
| cg06503255 | 5 | 140053350 | *DND1* | TSS200 | -4.97 | 6.77E-07 | 4.28E-07 |
| cg25984996 | 5 | 140098398 | *VTRNA1-2* | TSS200 | 5.18 | 2.19E-07 | 4.28E-07 |
| cg07539045 | 6 | 43245571 | *TTBK1* | Body | -5.33 | 9.85E-08 | 9.61E-07 |
| cg13305186 | 11 | 57508416 | *TMX2;TMX2;*<br>*C11orf31* | 3'UTR;Body;<br>TSS1500 | 4.95 | 7.56E-07 | 6.65E-08 |
| cg25744127 | 11 | 109292505 | *C11orf87* | TSS1500 | -4.99 | 6.12E-07 | 1.72E-07 |
| cg14258853 | 12 | 29935411 | *TMTC1* | 5'UTR | 5.18 | 2.25E-07 | 7.06E-08 |
| cg24305861 | 12 | 99564273 | *ANKS1B* | Body | -4.92 | 8.61E-07 | 1.28E-06 |
| cg09069446 | 16 | 4462122 | *CORO7* | Body | 4.93 | 8.21E-07 | 2.79E-07 |
| cg08460995 | 16 | 4526227 | *HMOX2;HMOX2;*<br>*NMRAL1* | TSS200;5'UTR;<br>TSS1500 | -5.03 | 4.87E-07 | 2.79E-07 |
| cg26619894 | 17 | 19249164 | *MIR1180;B9D1* | TSS1500;Body | 4.98 | 6.40E-07 | 7.77E-07 |
| cg22598241 | 19 | 2131519 | *AP3D1* | Body | -5.29 | 1.24E-07 | 1.22E-06 |
| cg04052466 | 19 | 2251061 | *AMH* | Body | 5.16 | 2.50E-07 | 1.22E-06 |
| cg09250473 | 19 | 50168907 | *IRF3;IRF3;*<br>*BCL2L12;BCL2L12* | 1stExon;5'UTR;<br>5'UTR;1stExon | -4.99 | 6.06E-07 | 2.19E-07 |
| cg13896476 | 22 | 31318546 | *MORC2-AS1* | Body | -6.28 | 3.37E-10 | 1.46E-05 |
| cg22416021 | 22 | 50177941 | *BRD1* | Body | 5.25 | 1.53E-07 | 3.31E-07 |
| cg26441486 | 22 | 50317300 | *CRELD2* | Body | 5.17 | 2.36E-07 | 3.31E-07 |

\* the strongest GWAS signals from SNPs within 1Mb of each gene

**Figure 4.**
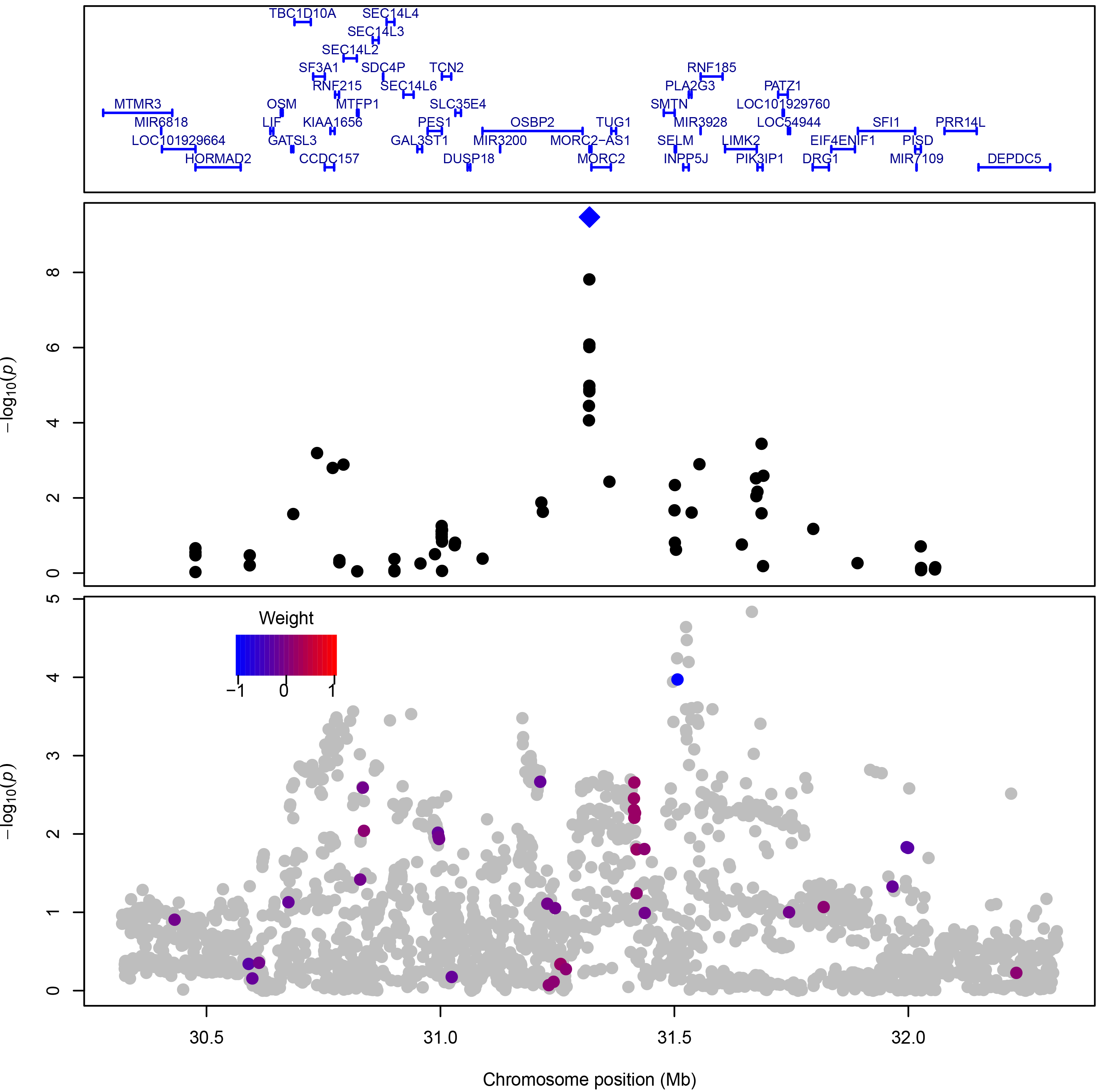
Regional association plot around the top significant CpG within *MORC1-AS.* The upper panel shows gene names and locations based on hg19 coordinates. The middle panel shows association strength of top significant CpG (blue diamond) and surrounding CpG sites (black) within 1Mb. The bottom panel shows association signals of SNPs from GWAS at the same region. SNPs are colored according to their weights contributing to the top significant CpG.

We identified 28 GWS CpGs with methylation-trait association evidence for BD (Figure 3b, Supplementary Table 2). Of these, 14 overlapped original GWAS risk loci and 14 were at least 1 Mb away from any GWS SNPs. Table 2 shows details and methylation-trait association statistics for the 10 CpG sites mapped to 11 unique genes. Notably, *CNIH2*, which encodes a subunit of the ionotropic glutamate receptor in the human brain, showed strong evidence for methylation-trait association at a CpG site in the promoter region (cg19026260, p=8.9×10^−9^), whereas the strongest GWAS signals within 1 Mb of *CNIH2* was only at p = 2.1×10^−7^. Figure 5 shows the regional association plot of methylation-trait associations around *CNIH2*, along with SNP association signals from GWAS and how their weights contribute to the top significant CpG.

**Table 2.**
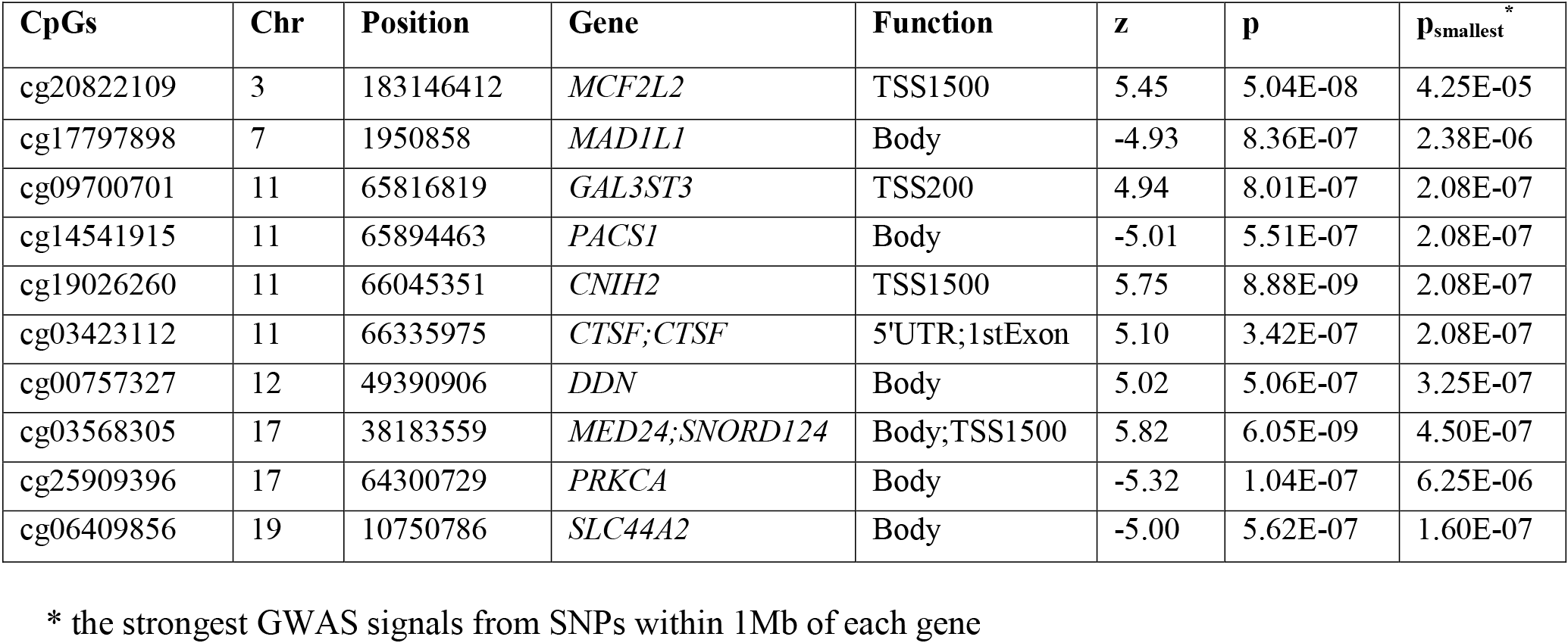
Information and methylation-trait association statistics for CpGs associated with bipolar disorder.

**Figure 5.**
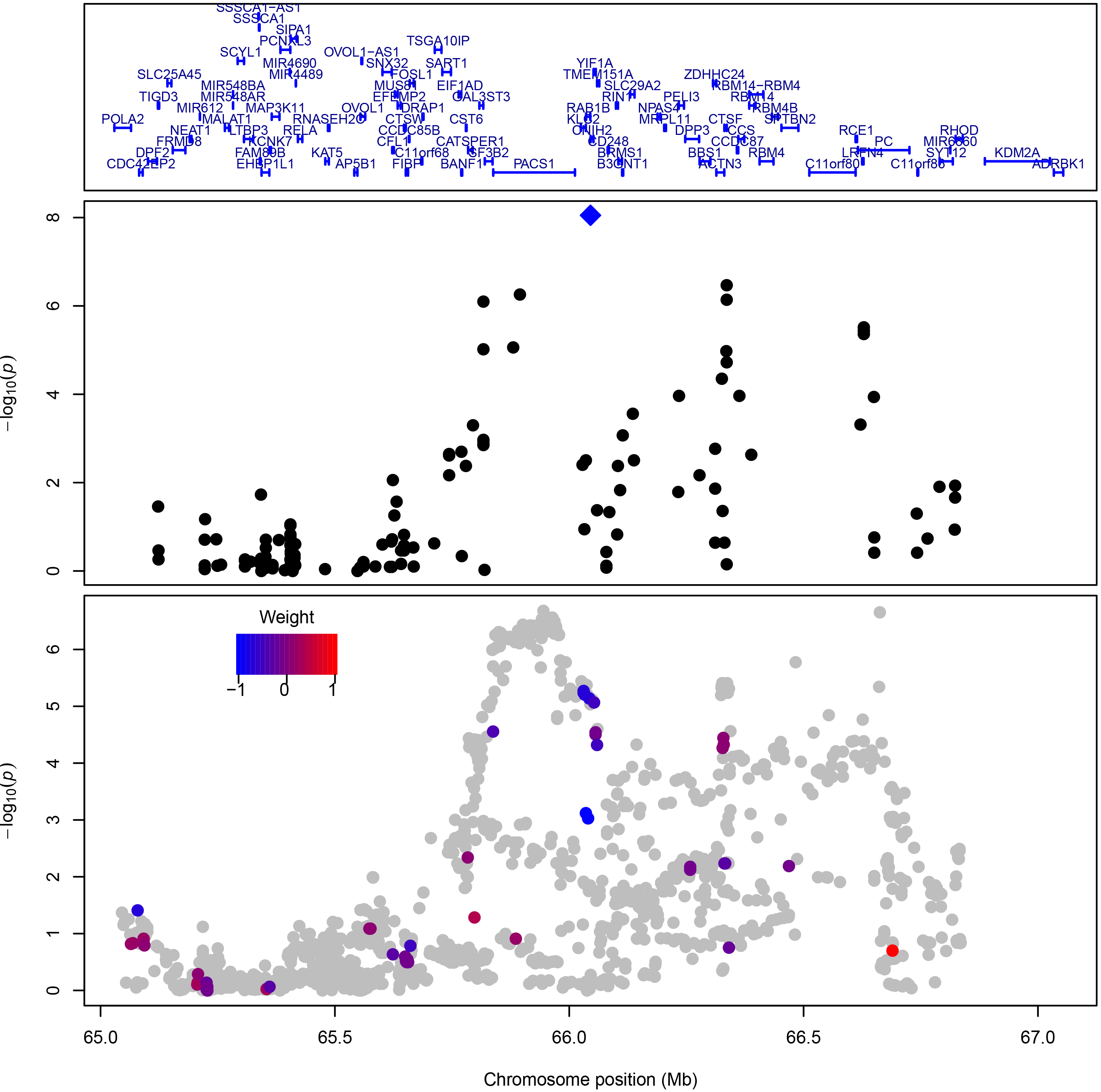
Regional association plot around the top significant CpG within *CNIH2.* The upper panel shows gene names and locations based on hg19 coordinates. The middle panel shows association strength of top significant CpG (blue diamond) and surrounding CpG sites (black) within 1Mb. The bottom panel shows association signals of SNPs from GWAS at the same region. SNPs are colored according to their weights contributing to the top significant CpG.

### Comparison with alternative analytical strategies

We evaluated three alternative strategies in their ability to detect the potential novel risk genes discovered above, which were at least 1 Mb away from any GWS SNPs (Supplementary Table 3). Among the 24 potential novel risk genes from PGC2 SCZ, only five genes were detected: *TMX2* (GATES, p = 6.9×10^−7^; VEGAS-sum, p = 1.0×10^−6^), *C11orf31* (VEGAS-sum, p = 1.0×10^−6^), *CORO7* (TWAS, p=2.3×10^−7^), *TMTC1* (TWAS, p = 1.8×10^−8^), and *BCL2L12* (VEGAS-sum, p = 2.0×10^−6^). For the ten potential novel risk genes from PGC2 BD, only *PACS1* was identified (VEGAS-sum, p=1.0×10^−6^).

### Gene set enrichment analysis

We conducted gene set enrichment analyses to examine whether genes with methylation-trait associations clustered into biological functional groups. Full gene set analysis results are shown in Supplementary table 4 (SCZ) and Supplementary table 5 (BD).

Figure 6 shows summary results for 60 gene sets with enriched methylation-trait association evidence for SCZ (q-values ≤ 0.05). We grouped gene sets into eight major clusters. The highlighted gene sets in each cluster include: 1) genes involved in synaptic transmission and plasticity (dark red); 2) RBFOX1, RBFOX2, RBFOX3 regulatory networks, FMRP targets, synaptic genes, voltage-gated calcium channel transporters, missense constrained genes, mutation intolerant genes; SCZ candidate genes (green); 3) genes encoding voltage-gated calcium channels, genes related to membrane depolarization during action potential, genes associated with SCZ (p <10^−4^), autism *de novo* genes (orange); 4) microRNA targets, SCZ *de novo* genes (blue); 5) microRNA targets, chromatin regulator CHD8 targets (red); 6) genes involved in antigen processing and presentation, genes associated with BP (p <10^−4^) (brown); 7) genes involved in cell adhesion (black); 8) genes involved in axoneme assembly (dark blue).

**Figure 6.**
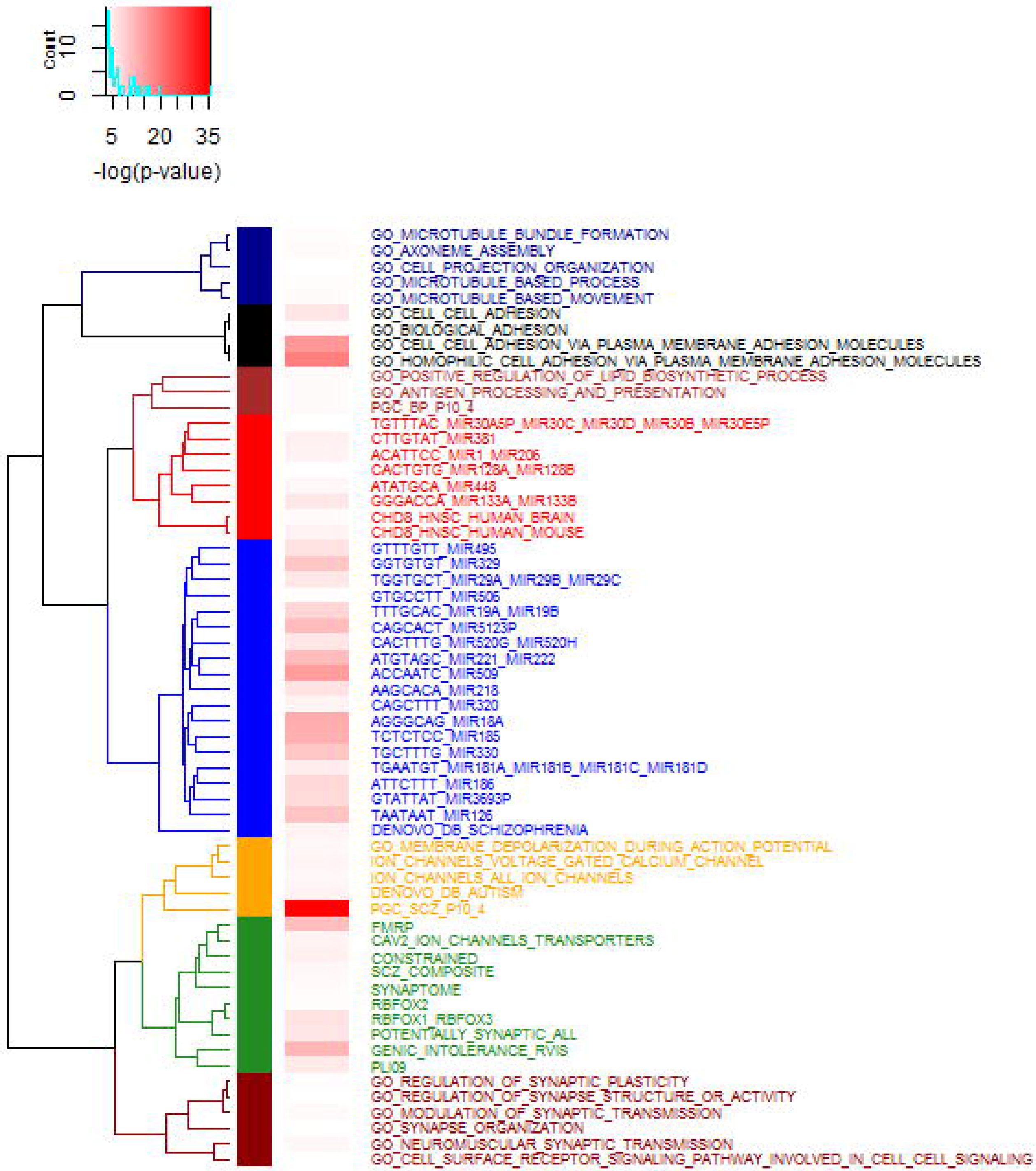
Hierarchical clustering of significant gene sets in schizophrenia. Gene sets were grouped into eight clusters as indicated by different colors. The color gradient indicates the enrichment p-values of each gene set as shown in the middle panel.

Supplementary Figure 3 shows summary results for 51 gene sets with enriched methylation-trait association evidence for BD (q-values ≤ 0.05). Gene sets were grouped into five major clusters. The highlighted gene sets in each cluster include: 1) microRNA targets, SCZ *de novo* genes, and genes involved in cell adhesion (blue); 2) RBFOX1, RBFOX2, RBFOX3 regulatory networks, FMRP targets, synaptome, mutation intolerant genes (red); 3) genes involved in synaptic plasticity and transmission (green); 4) ion channel genes, missense constrained genes, calcium signaling pathway (purple); 5) genes involved in integrin signaling, genes associated with BP and depression (p <10^−4^) (orange).

## Discussion

In the current study, we have shown that DNAm levels are heritable for ~17% of the CpG sites examined in human prefrontal cortex. Heritable DNAm sites tended to be more variable and enriched in intergenic and regulatory regions in brain. Through imputation of methylation-trait association from GWAS summary statistics, we identified known and potentially novel risk genes with methylation-trait association evidence that were not detectable using three alternative strategies. Gene set enrichment analysis for genes with methylation-trait association evidence revealed pathways related to neuronal functions, as well as other biological mechanisms potentially underlying psychiatric disorders, consistent with prior analyses based on GWAS (46) and gene expression data (47–49).

Heritability estimates of DNAm levels may vary across tissues and study designs. Twin studies reported an average heritability of 12–18% in whole blood (7, 50), 5% in placenta (7), and 19% in adipose tissue (28). A family study showed an average heritability of 13% in CD4^+^ cells (3). One study reported an average SNP-heritability of 29% across heritable CpG sites in colorectum (51). There has been to date only one study that estimated the SNP-heritability of DNAm in brain, but that was limited to ~27,000 CpGs located primarily in promoters (8). We found modest correlation of heritability estimates (r = 0.48) for a common subset of CpGs between our study and the frontal cortex results of Quon *et al* (8). The variation of heritability estimates may be attributable to differences in brain samples and analytical approach. For example, we used a more narrowly defined single prefrontal cortical region---dorsolateral prefrontal cortex---from samples with a wide range of ages, whereas Quon *et al* used the whole frontal cortex largely from adults; we used SNPs within 1 Mb window of CpGs, but Quon *et al* used a window size of 50 kb. Importantly, our study differs from the previous one by virtue of a more than tenfold increase of CpG sites (~380,000) across a much broader portion of the genome, including intergenic and gene body regions. In addition, our study has a larger sample size than the previous one (238 versus 150) and thus higher power to detect heritable CpG sites. We found a higher mean heritability for CpG sites in intergenic regions than for those at promoter regions. We also observed that heritable CpG sites tended to be located in intergenic regulatory genomic regions, suggesting a potentially important role in regulation of gene expression.

We evaluated three modeling schemes for their ability to predict the cis-genetic component of DNAm levels for CpG sites with a nominal significance level for heritability estimates. We found the highest performance for elastic net and the lowest performance for the top SNP method, suggesting that DNAm levels tend to be influenced by multiple SNPs. This observation is consistent with the result from a previous study in which elastic net achieved the best performance for predicting the cis-genetic component of gene expression using SNPs within 1 Mb of a gene (52). Future work might employ other approaches to improve the prediction accuracy for DNAm levels. For example, recent work has included imputation of gene expression using the best linear predictor and the Bayesian linear mixed model (38), which can also be evaluated for DNAm prediction in the future.

Compared to a single marker association, the proposed MWAS features similar advantages as the TWAS method that aims to impute expression-trait association from GWAS summary statistics. First, both methods have significantly reduced multiple testing burdens compared to GWAS, though there are slightly more DNAm sites tested in MWAS than genes tested in TWAS. Second, because MWAS aims to detect methylation differences that are genetically driven, significant findings from MWAS may indicate a causal relationship between differential methylation and trait, whereas for EWAS the identified methylation difference may be a consequence of disease. This was further supported by the limited correlation (r = 0.025) between the MWAS signals (Z-score) from PGC SCZ2 GWAS and the EWAS signals (T-score) from the comparison of brains of SCZ versus controls identified in our previous study (13). Third, MWAS may increase the power to detect DNAm sites of small effect by taking advantage of GWAS summary statistics from large samples. In addition, MWAS has a unique advantage of higher coverage of genomic regions compared to TWAS. For example, MWAS covers CpGs in more than 10,000 genes in our study, whereas TWAS only contains ~5,000 genes for the prefrontal cortex region examined. Furthermore, DNAm levels may be a more proximal read out of genetic variation than gene expression levels, at least as measured in homogenate tissue at one time point (53), and MWAS may have greater power than TWAS in risk gene discovery in some situations.

We identified known and novel risk genes by applying MWAS to two major psychiatric disorders. We detected a number of potentially novel risk genes through application of MWAS to PGC2 SCZ GWAS (table 1). For example, *KCNN3* was suspected to be involved in SCZ by candidate gene and gene expression studies (54–56); *ANKS1B* was among the top findings in a genome-wide pharmacogenomic study of antipsychotic treatment response in SCZ, which mediates the effect of quetiapine on working memory (57); *TTBK1* was found to harbor *de novo* mutations in sporadic cases of childhood onset SCZ (58); earlier linkage and candidate gene analysis showed the association evidence of *BRD1* with SCZ (59), which was further supported by a recent functional study, demonstrating that *BRD1* regulates behavior, neurotransmission, and expression of SCZ risk-enriched gene sets in mice (60). The association between *CNIH2* and BD is also noteworthy. *CNIH2* encodes a subunit of the ionotropic glutamate receptor in the human brain, and has been shown to influence the efficacy of excitatory synaptic transmission (61).

Gene set enrichment analysis of MWAS association signals not only revealed pathways related to neuronal functions, such as synaptic genes and FMRP targets, but also highlighted other biological mechanisms that may underlie psychiatric disorders. One implication is the likely involvement of chromatin remodeling in SCZ as indicated by the enrichment of association signals in CHD8 targets.

CHD8 is a DNA binding protein and acts as a chromatin remodeling factor. Genes targeted by CHD8 in human midfetal brain were enriched for functions related to transcriptional regulation and chromatin modification (62). *CHD8* is also one of the most frequently mutated genes in autism (63–65), consistent with other data suggesting some genetic overlap between autism and SCZ (66). Perhaps the most notable highlight from gene set analysis is the involvement of microRNA-related regulation mechanism in psychiatric disorders. We found significant enrichment of microRNA targets for both disorders. Interestingly, targets of the hsa-miR-221 and hsa-miR-222 were found to be enriched for both disorders. Intriguingly, reduced expression of miR-221 was observed following chronic treatment with the antidepressant paroxetine in both rat hippocampus (67) and human lymphoblastoid cell lines (68). Additionally, we found significant enrichment of association signals in mutation-intolerant genes or missense constrained genes in both disorders. Our results are consistent with recent studies that have shown an enrichment of *de novo* mutations or rare risk variants in mutation-intolerant genes for neurodevelopmental disorders such as autism, intellectual disability and developmental delay, as well as SCZ (69–72).

This work should be viewed in light of several limitations. First, MWAS was designed to identify methylation-trait associations for DNAm sites that have a genetic component. It is therefore not suited to detect methylation differences caused exclusively by environmental factors. That being said, our heritability estimates themselves may be diluted by environmental factors. Indeed, the average heritability estimate from the overall samples was slightly lower than the estimate from the neurotypical samples alone, consistent with the possibility that systematic environmental factors (e.g. medical treatment) diminish heritability statistics in patients. Second, the number of DNAm sites that can be accurately imputed is limited by the training sample size and tissue types. Future studies will benefit from larger training samples and diverse tissues related to disease. Third, the Illumina CpG chip that we used is a relatively low resolution survey of the DNA methylation landscape, and though in includes many CpGs not in the 5’ domain of genes and in CpG islands, the majority of CpGs are still in these regions. Fourth, MWAS can only identify risk SNPs that act through regulation of DNAm levels; it is not able to detect SNPs acting through mechanisms that are independent of DNAm regulation. Therefore, we view MWAS as by no means a replacement for current methods, such as TWAS or other gene-based tests, but as a complementary approach for uncovering genetic underpinnings underlying complex diseases.

In summary, we have shown the heritability pattern of DNAm in human prefrontal cortex. We further demonstrated the power of integrating the brain methylome with GWAS for psychiatric risk gene discovery, an approach which has potential applications in other brain-related disorders or traits.

## Acknowledgments

The computations work for this study was supported by National Institutes of Health grants R01 AA022994 and AA024486 (to S.H.). All of the human brain data was generated from the Lieber Institute for Brain Development and supported by Institute funds. We are grateful for the vision and generosity of the Lieber and Maltz Families who made this work possible. We thank the families who donated to this research. We acknowledge the Psychiatric Genomics Consortium for making the GWAS results publicly available.

---

**Supplementary Figure 1.**
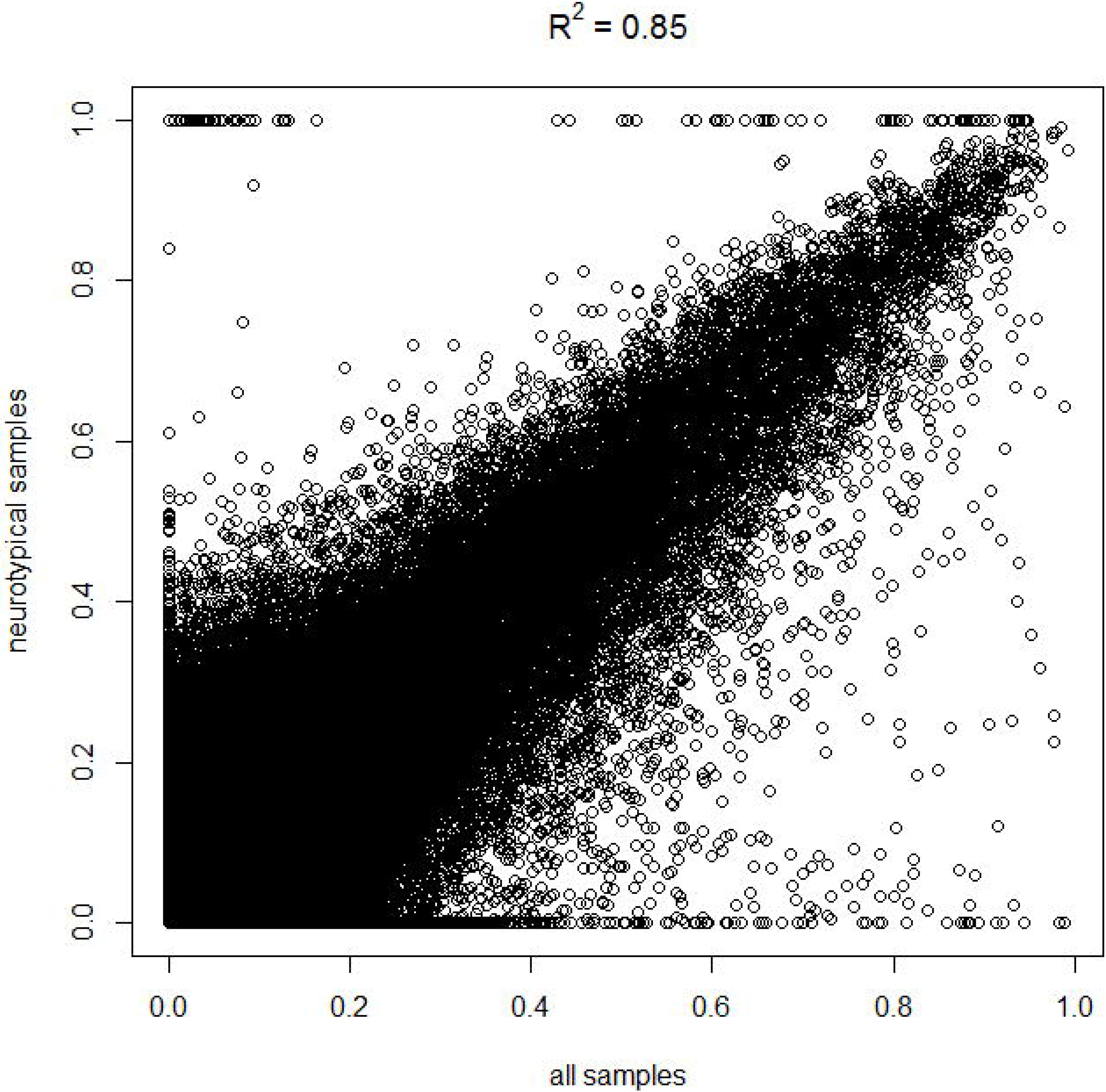
Scatter plot for heritability estimates from the overall samples and neurotypical samples.

**Supplementary Figure 2.**
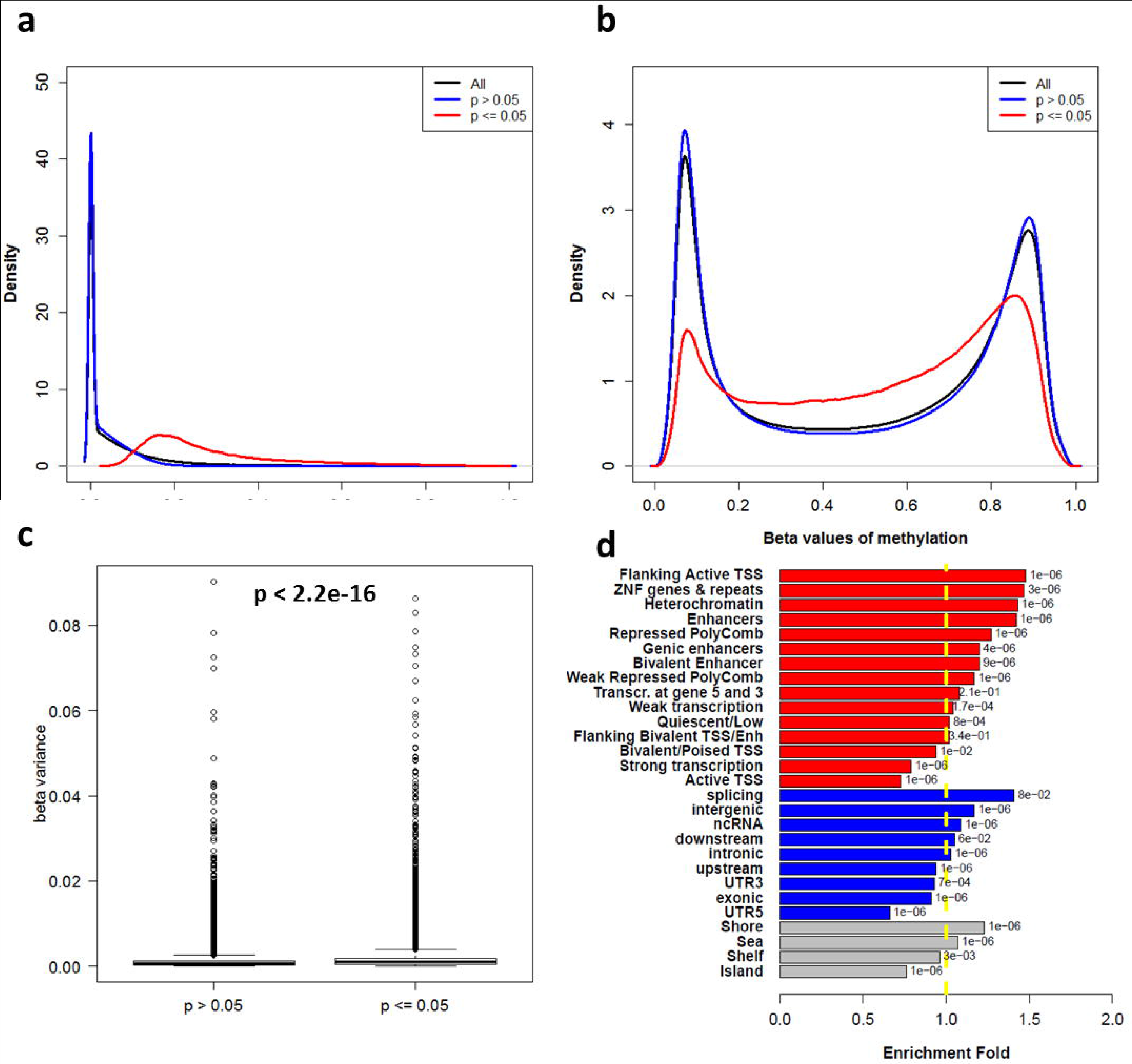
SNP-based heritability analysis of DNAm levels from neurotypical samples: a) Distribution of heritability estimates for all, heritable (p ≤ 0.05), and non-heritable (p > 0.05) CpG sites; b) Distribution of methylation levels (beta-value) for all, heritable (p ≤ 0.05), and non-heritable (p > 0.05) CpG sites; c) Boxplot for variance of methylation levels (beta-value) for heritable (p ≤ 0.05) and non-heritable (p > 0.05) CpG sites; d) Enrichment of heritable CpG sites across genomic features within three different contexts: genomic locations (blue), distance to CpG islands (grey), and functional states in the dorsolateral prefrontal cortex (red). The x-axis represents enrichment fold. The y-axis labels the different features examined. The dotted yellow line indicates no enrichment (fold = 1). The numbers next to each bar are enrichment p-values.

**Supplementary Figure 3.**
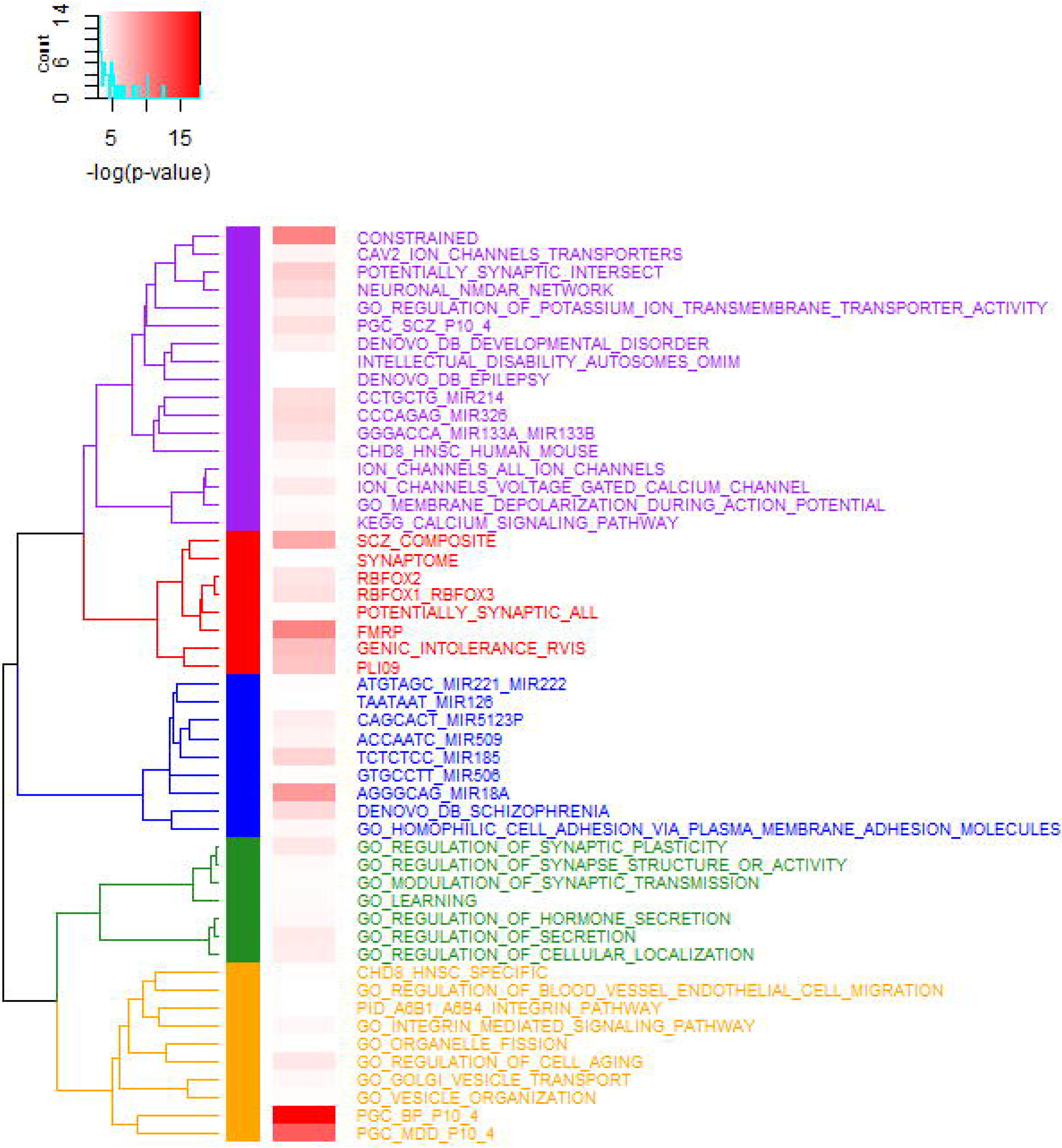
Hierarchical clustering of significant gene sets in bipolar disorder. Gene sets were grouped into five clusters as indicated by different colors. The color gradient indicates the enrichment p-values of each gene set as shown in the middle panel.

